# #COVIDisAirborne: AI-Enabled Multiscale Computational Microscopy of Delta SARS-CoV-2 in a Respiratory Aerosol

**DOI:** 10.1101/2021.11.12.468428

**Authors:** Abigail Dommer, Lorenzo Casalino, Fiona Kearns, Mia Rosenfeld, Nicholas Wauer, Surl-Hee Ahn, John Russo, Sofia Oliveira, Clare Morris, Anthony Bogetti, Anda Trifan, Alexander Brace, Terra Sztain, Austin Clyde, Heng Ma, Chakra Chennubhotla, Hyungro Lee, Matteo Turilli, Syma Khalid, Teresa Tamayo-Mendoza, Matthew Welborn, Anders Christensen, Daniel G. A. Smith, Zhuoran Qiao, Sai Krishna Sirumalla, Michael O’Connor, Frederick Manby, Anima Anandkumar, David Hardy, James Phillips, Abraham Stern, Josh Romero, David Clark, Mitchell Dorrell, Tom Maiden, Lei Huang, John McCalpin, Christopher Woods, Alan Gray, Matt Williams, Bryan Barker, Harinda Rajapaksha, Richard Pitts, Tom Gibbs, John Stone, Daniel Zuckerman, Adrian Mulholland, Thomas Miller, Shantenu Jha, Arvind Ramanathan, Lillian Chong, Rommie Amaro

**Affiliations:** UC San Diego; Oregon Health & Science University; University of Bristol; University of Pittsburgh; Argonne National Laboratory; University of Illinois at Urbana-Champaign; University of Chicago; Freie Universität Berlin; Brookhaven National Lab & Rutgers University; University of Oxford; Entos, Inc.; California Institute of Technology; NVIDIA Corporation; Pittsburgh Supercomputing Center; Texas Advanced Computing Center; Oracle for Research

**Keywords:** molecular dynamics, deep learning, multiscale simulation, weighted ensemble, computational virology, SARS-CoV-2, aerosols, COVID-19, HPC, AI, GPU, Delta

## Abstract

We seek to completely revise current models of airborne transmission of respiratory viruses by providing never-before-seen atomic-level views of the SARS-CoV-2 virus within a respiratory aerosol. Our work dramatically extends the capabilities of multiscale computational microscopy to address the significant gaps that exist in current experimental methods, which are limited in their ability to interrogate aerosols at the atomic/molecular level and thus obscure our understanding of airborne transmission. We demonstrate how our integrated data-driven platform provides a new way of exploring the composition, structure, and dynamics of aerosols and aerosolized viruses, while driving simulation method development along several important axes. We present a series of initial scientific discoveries for the SARS-CoV-2 Delta variant, noting that the full scientific impact of this work has yet to be realized.

**ACM Reference Format:** Abigail Dommer^1†^, Lorenzo Casalino^1†^, Fiona Kearns^1†^, Mia Rosenfeld^1^, Nicholas Wauer^1^, Surl-Hee Ahn^1^, John Russo,^2^ Sofia Oliveira^3^, Clare Morris^1^, AnthonyBogetti^4^, AndaTrifan^5,6^, Alexander Brace^5,7^, Terra Sztain^1,8^, Austin Clyde^5,7^, Heng Ma^5^, Chakra Chennubhotla^4^, Hyungro Lee^9^, Matteo Turilli^9^, Syma Khalid^10^, Teresa Tamayo-Mendoza^11^, Matthew Welborn^11^, Anders Christensen^11^, Daniel G. A. Smith^11^, Zhuoran Qiao^12^, Sai Krishna Sirumalla^11^, Michael O’Connor^11^, Frederick Manby^11^, Anima Anandkumar^12,13^, David Hardy^6^, James Phillips^6^, Abraham Stern^13^, Josh Romero^13^, David Clark^13^, Mitchell Dorrell^14^, Tom Maiden^14^, Lei Huang^15^, John McCalpin^15^, Christo- pherWoods^3^, Alan Gray^13^, Matt Williams^3^, Bryan Barker^16^, Harinda Rajapaksha^16^, Richard Pitts^16^, Tom Gibbs^13^, John Stone^6^, Daniel Zuckerman^2^*, Adrian Mulholland^3^*, Thomas MillerIII^11,12^*, ShantenuJha^9^*, Arvind Ramanathan^5^*, Lillian Chong^4^*, Rommie Amaro^1^*. 2021. #COVIDisAirborne: AI-Enabled Multiscale Computational Microscopy ofDeltaSARS-CoV-2 in a Respiratory Aerosol. In *Supercomputing ‘21: International Conference for High Performance Computing, Networking, Storage, and Analysis*. ACM, New York, NY, USA, 14 pages. https://doi.org/finalDOI

## 1 JUSTIFICATION

We develop a novel HPC-enabled multiscale research framework to study aerosolized viruses and the full complexity of species that comprise them. We present technological and methodological advances that bridge time and length scales from electronic structure through whole aerosol particle morphology and dynamics.

## 2 PERFORMANCE ATTRIBUTES

| Performance Attribute | Our Submission |
| --- | --- |
| Category of achievement | Scalability, Time-to-solution |
| Type of method used | Explicit, Deep Learning |
| Results reported on the basis of | Whole application including I/O |
| Precision reported | Mixed Precision |
| System scale | Measured on full system |
| Measurement mechanism | Hardware performance counters, Application timers, Performance Modeling |

## 3 OVERVIEW OF THE PROBLEM

Respiratory pathogens, such as SARS-CoV-2 and influenza, are the cause of significant morbidity and mortality worldwide. These respiratory pathogens are spread by virus-laden aerosols and droplets that are produced in an infected person, exhaled, and transported through the environment (Wang et al., 2021) (Fig 1). Medical dogma has long focused on droplets as the main transmission route for respiratory viruses, where either a person has contact with an infected surface (fomites) or direct droplet transmission by close contact with an infected individual. However, as we continue to observe with SARS-CoV-2, airborne transmission also plays a significant role in spreading disease. We know this from various super spreader events, e.g., during a choir rehearsal (Miller et al., 2021). Intervention and mitigation decisions, such as the relative importance of surface cleaning or whether and when to wear a mask, have unfortunately hinged on a weak understanding of aerosol transmission, to the detriment of public health.

**Figure 1:**
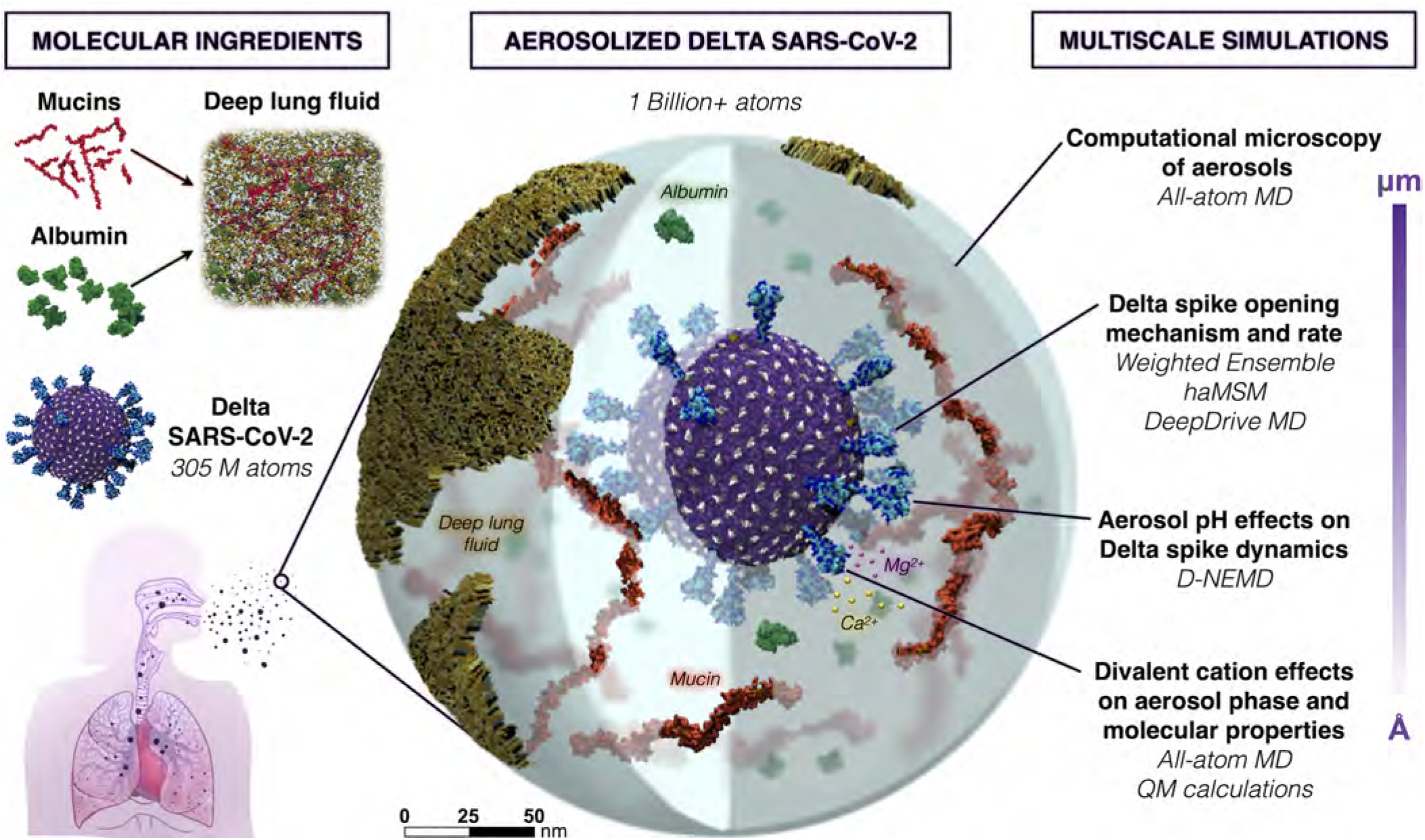
Overall schematic depicting the construction and multiscale simulations of Delta SARS-CoV-2 in a respiratory aerosol. (N.B.: The size of divalent cations has been increased for visibility.)

A central challenge to understanding airborne transmission has been the inability of experimental science to reliably probe the structure and dynamics of viruses once they are inside respiratory aerosol particles. Single particle experimental methods have poor resolution for smaller particles (<1 micron) and are prone to sample destruction during collection. Airborne viruses are present in low concentrations in the air and are similarly prone to viral inactivation during sampling. In addition, studies of the initial infection event, for example in the deep lung, are limited in their ability to provide a detailed understanding of the myriad of molecular interactions and dynamics taking place *in situ*. Altogether, these knowledge gaps hamper our collective ability to understand mechanisms of infection and develop novel effective antivirals, as well as prevent us from developing concrete, science-driven mitigation measures (e.g., masking and ventilation protocols).

Here, we aim to reconceptualize current models of airborne transmission of respiratory viruses by providing never-before-seen views of viruses within aerosols. Our approach relies on the use of all-atom molecular dynamics (MD) simulations as a multiscale ‘computational microscope’ MD simulations can synthesize multiple types of biological data (e.g., multiresolution structural datasets, glycomics, lipidomics, etc.) into cohesive, biologically ‘accurate’ structural models. Once created, we then approximate the model down to its many atoms, creating trajectories of its time dependent dynamics under cell-like (or in this case, aerosol-like) conditions. Critically, MD simulations are more than just ‘pretty movies.’ MD equations are solved in a theoretically rigorous manner, allowing us to compute experimentally testable macroscopic observables from time-averaged microscopic properties. What this means is that we can directly connect MD simulations with experiments, each validating and providing testable hypotheses to the other, which is the real power of the approach. An ongoing challenge to the successful application of such methods, however, is the need for technological and methodological advances that make it possible to access length scales relevant to the study of large, biologically complex systems (spanning nanometers to ~one micron in size) and, correspondingly, longer timescales (microseconds to seconds).

Such challenges and opportunities manifest in the study of aerosolized viruses. Aerosols are generally defined as being less than 5 microns in diameter, able to float in the air for hours, travel significant distances (e.g., can fill a room, like cigarette smoke), and be inhaled. Fine aerosols < 1 micron in size can stay in the air for over 12 hours and are enriched with viral particles (Coleman et al.,2021, Fennelly, 2020). Our work focuses on these finer aerosols that travel deeper into the respiratory tract. Several studies provide the molecular recipes necessary to reconstitute respiratory aerosols according to their actual biologically-relevant composition (Vejerano and Marr, 2018, Walker et al., 2021). These aerosols can contain lipids, cholesterol, albumin (protein), various mono- and di-valent salts, mucins, other surfactants, and water (Fig 1). Simulations of aerosolized viruses embody a novel framework for the study of aerosols: they will allow us and others to tune different species, relative humidity, ion concentrations, etc. to match experiments that can directly and indirectly connect to and inform our simulations, as well as test hypotheses. Some of the species under study here, e.g., mucins, have not yet been structurally characterized or explored with simulations and thus the models we generate are expected to have impact beyond their roles in aerosols.

In addition to varying aerosol composition and size, the viruses themselves can be modified to reflect new variants of concern, where such mutations may affect interactions with particular species in the aerosol that might affect its structural dynamics and/or viability. The virion developed here is the Delta variant (B.1.617.2 lineage) of SARS-CoV-2 (Fig 2), which presents a careful integration of multiple biological datasets: (1) a complete viral envelope with realistic membrane composition, (2) fully glycosylated full-length spike proteins integrating 3D structural coordinates from multiple cryoelectron microscopy (cryoEM) studies (Bangaru et al., 2020,McCallum et al., 2021, Walls et al., 2020, Wrapp et al., 2020) (3) all biologically known features (post-translational modifications, palmitoylation, etc.), (4) any other known membrane proteins (e.g., the envelope (E) and membrane (M) proteins), and (5) virion size and patterning taken directly from cryoelectron tomography (cryoET). Each of the individual components of the virus are built up before being integrated into the composite virion, and thus represent useful molecular-scale scientific contributions in their own right (Casalino et al., 2020, Sztain et al., 2021).

**Figure 2:**
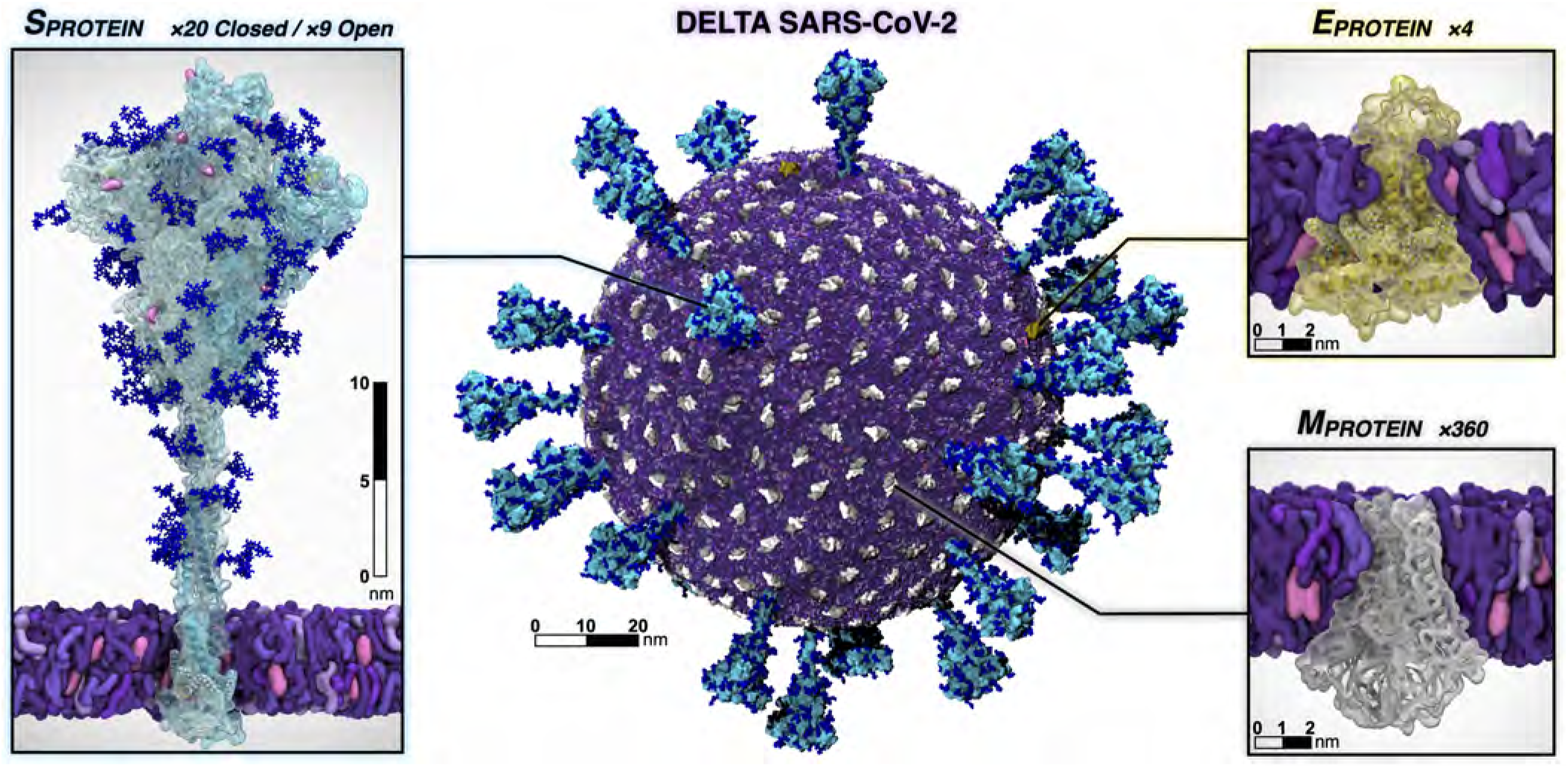
Individual protein components of the SARS-CoV-2 Delta virion. The spike is shown with the surface in cyan and with Delta’s mutated residues and deletion sites highlighted in pink and yellow, respectively. Glycans attached to the spike are shown in blue. The E protein is shown in yellow and the M protein is shown in silver and white. Visualized with VMD.

Altogether in this work, we dramatically extend the capabilities of data-driven, multiscale computational microscopy to provide a new way of exploring the composition, structure, and dynamics of respiratory aerosols. While a seemingly limitless number of putative hypotheses could result from these investigations, the first set of questions we expect to answer are: *How does the virus exist within a droplet of the same order of magnitude in size, without being affected by the air-water interface, which is known to destroy molecular structure (D’Imprima et al., 2019)? How does the biochemical composition of the droplet, including pH, affect the structural dynamics of the virus? Are there species within the aerosols that “buffer” the viral structure from damage, and are there particular conditions under which the impact of those species changes?* Our simulations can also provide specific parameters that can be included in physical models of aerosols, which still assume a simple water or water-salt composition even though it is well known that such models, e.g., using kappa-Kohler theory, break down significantly as the molecular species diversify (Petters and Kreidenweis, 2007).

## 4 CURRENT STATE OF THE ART

Current experimental methods are unable to directly interrogate the atomic-level structure and dynamics of viruses and other molecules within aerosols. Here we showcase computational microscopy as a powerful tool capable to overcome these significant experimental limitations. We present the major elements of our multiscale computational microscope and how they come together in an integrated manner to enable the study of aerosols across multiple scales of resolution. We demonstrate the impact such methods can bring to bear on scientific challenges that until now have been intractable, and present a series of new scientific discoveries for SARS-CoV-2.

### 4.1 Parallel molecular dynamics

All-atom molecular dynamics simulation has emerged as an increasingly powerful tool for understanding the molecular mechanisms underlying biophysical behaviors in complex systems. Leading simulation engines, NAMD (Phillips et al., 2020), AMBER (Case et al.), and GROMACS (Páll et al., 2020), are broadly useful, with each providing unique strengths in terms of specific methods or capabilities as required to address a particular biological question, and in terms of their support for particular HPC hardware platforms. Within the multiscale computational microscopy platform developed here, we show how each of these different codes contributes different elements to the overall framework, oftentimes utilizing different computing modalities/architectures, while simultaneously extending on state-of-the-art for each. Structure building, simulation preparation, visualization, and post-hoc trajectory analysis are performed using VMD on both local workstations and remote HPC resources, enabling modeling of the molecular systems studied herein (Humphrey et al., 1996, Sener et al., 2021, Stone et al., 2013a,b,2016b). We show how further development of each of these codes, considered together within the larger-scale collective framework, enables the study of SARS-CoV-2 in a wholly novel manner, with extension to numerous other complex systems and diseases.

### 4.2 AI-enhanced WE simulations

Because the virulence of the Delta variant of SARS-CoV-2 may be partly attributable to spike protein (S) opening, it is of pressing interest to characterize the mechanism and kinetics of the process. Although S-opening in principle can be studied via conventional MD simulations, in practice the system complexity and timescales make this wholly intractable. Splitting strategies that periodically replicate promising MD trajectories, among them the weighted ensemble (WE) method (Huber and Kim, 1996, Zuckerman and Chong, 2017), have enabled simulations of the spike opening of WT SARS-CoV-2 (Sztain et al., 2021, Zimmerman et al., 2021). WE simulations can be orders of magnitude more efficient than conventional MD in generating pathways and rate constants for rare events (e.g., protein folding (Adhikari et al., 2019) and binding (Saglam and Chong, 2019)). The WESTPA software for running WE (Zwier et al.,2015) is well-suited for high-performance computing with nearly perfect CPU/GPU scaling. The software is interoperable with any dynamics engine, including the GPU-accelerated AMBER dynamics engine (Salomon-Ferrer et al., 2013) that is used here. As shown below, major upgrades to WESTPA (v. 2.0) have enabled a dramatic demonstration of spike opening in the Delta variant (Figs. 5, 6) and exponentially improved analysis of spike-opening kinetics.

The integration of AI techniques with WE can further enhance the efficiency of sampling rare events (Brace et al., 2021b, Casalino et al., 2021, Noé, 2020). One frontier area couples unsupervised linear and non-linear dimensionality reduction methods to identify collective variables/progress coordinates in high-dimensional molecular systems (Bhowmik et al., 2018, Clyde et al., 2021). Such methods may be well suited for analyzing the aerosolized virus. Integrating these approaches with WE simulations is advantageous in sampling the closed → open transitions in the Delta S landscape (Fig. 5) as these unsupervised AI approaches automatically stratify progress coordinates (Fig. 5D).

### 4.3 Dynamical Non-Equilibrium MD

Aerosols rapidly acidify during flight via reactive uptake of atmospheric gases, which is likely to impact the opening/closing of the S protein (Vejerano and Marr, 2018, Warwicker, 2021). Here, we describe the extension of dynamical non-equilibrium MD (D-NEMD) (Ciccotti and Ferrario, 2016) to investigate pH effects on the Delta S. D-NEMD simulations (Ciccotti and Ferrario, 2016) are emerging as a useful technique for identifying allosteric effects and communication pathways in proteins (Galdadas et al., 2021, Oliveira et al., 2019), including recently identifying effects of linoleic acid in the WT spike (Oliveira et al., 2021b). This approach complements equilibrium MD simulations, which provide a distribution of configurations as starting points for an ensemble of short non-equilibrium trajectories under the effect of the external perturbation. The response of the protein to the perturbation introduced can then be determined using the Kubo-Onsager relation (Ciccotti and Ferrario,2016, Oliveira et al., 2021a) by directly tracking the change in atomic positions between the equilibrium and non-equilibrium simulations at equivalent points in time (Oliveira et al., 2021a).

### 4.4 OrbNet

Ca^2+^ ions are known to play a key role in mucin aggregation in epithelial tissues (Hughes et al., 2019). Our RAV simulations would be an ideal case-study to probe such complex interactions between Ca^2+^, mucins, and the SARS-CoV-2 virion in aerosols. However, Ca^2+^ binding energies can be difficult to capture accurately due to electronic dispersion and polarization, terms which are not typically modeled in classical mechanical force fields. Quantum mechanical (QM) methods are uniquely suited to capture these subtle interactions. Thus, we set out to estimate the correlation in Ca^2+^ binding energies between CHARMM36m and quantum mechanical estimates enabled via AI with OrbNet. Calculation of energies with sufficient accuracy in biological systems can, in many cases, be adequately described with density functional theory (DFT). However, its high cost limits the applicability of DFT in comparison to fixed charge force-fields. To capture quantum quality energetics at a fraction of the computational expense, we employ a novel approach (OrbNet) based on the featurization of molecules in terms of symmetry-adapted atomic orbitals and the use of graph neural network methods for deep-learning quantum-mechanical properties (Qiao et al., 2020). Our method outperforms existing methods in terms of its training efficiency and transferable accuracy across diverse molecular systems, opening a new pathway for replacing DFT in large-scale scientific applications such as those explored here. (Christensen et al., 2021).

## 5 INNOVATIONS REALIZED

### 5.1 Construction and simulation of SARS-CoV-2 in a respiratory aerosol

Our approach to simulating the entire aerosol follows a composite framework wherein each of the individual molecular pieces is refined and simulated on its own before it is incorporated into the composite model. Simulations of each of the components are useful in their own right, and often serve as the basis for biochemical and biophysical validation and experiments (Casalino et al., 2020).

Throughout, we refer to the original circulating SARS-CoV-2 strain as “WT”, whereas all SARS-CoV-2 proteins constructed in this work represent the Delta variant (Fig. 2). All simulated membranes reflect mammalian ER-Golgi intermediate compartment (ERGIC) mimetic lipid compositions. VMD (Humphrey et al., 1996,Stone et al., 2016a), psfgen (Phillips et al., 2005), and CHARMM-GUI (Park et al., 2019) were used for construction and parameterization. Topologies and parameters for simulations were taken from CHARMM36m all-atom additive force fields (Beglov and Roux, 1994,Guvench et al., 2009, Han et al., 2018, Huang and Mackerell, 2013,Huang et al., 2017, Klauda et al., 2010,Venable et al., 2013). NAMD was used to perform MD simulations (Phillips et al., 2020), adopting similar settings and protocols as in (Casalino et al., 2020). All systems underwent solvation, charge neutralization, minimization, heating and equilibration prior to production runs. Refer to Table 1 for Abbreviations, PBC dimensions, total number of atoms, and total equilibration times for each system of interest.

**Table 1:** Summary of all systems constructed in this work. See Fig 3 for illustration of aerosol construction.

| <sup>a</sup>systems | <sup>b</sup>Abb. | <sup>c</sup>(Å × Å × Å) | <sup>d</sup>N<sub>a</sub> | <sup>e</sup>(ns) |
| --- | --- | --- | --- | --- |
| <sup>f</sup>M dimers | M | 125 × 125 × 124 | 164,741 | 700 |
| <sup>f</sup>E pentamers | E | 123 × 125 × 102 | 136,775 | 41 |
| **Spikes:** |  |  |  |  |
| <sup>f</sup>(open) | S | 206 × 200 × 410 | 1,692,444 | 330 |
| <sup>f</sup>(closed) | S | 204 × 202 × 400 | 1,658,224 | 330 |
| <sup>g</sup>(closed, head) | SH | 172 × 184 × 206 | 615,593 | 73μs |
| **Mucins:** |  |  |  |  |
| <sup>f</sup>short mucin 1 | m<sub>1</sub> | 123 × 104 × 72 | 87,076 | 25 |
| <sup>f</sup>short mucin 2 | m<sub>2</sub> | 120 × 101 × 72 | 82,155 | 25 |
| <sup>f</sup>long mucin 1 | m<sub>3</sub> | 810 × 104 × 115 | 931,778 | 23 |
| <sup>f</sup>long mucin 2 | m<sub>4</sub> | 904 × 106 × 109 | 997,029 | 15 |
| <sup>f</sup>long mucin 3 | m<sub>5</sub> | 860 × 111 × 113 | 1,040,215 | 18 |
| <sup>f</sup>S+m1/m2+ALB | SMA | 227 × 229 × 433 | 2,156,689 | 840 |
| <sup>f</sup>Virion | V | 1460 × 1460 × 1460 | 305,326,834 | 41 |
| <sup>f</sup>Resp.Aero.+Vir. | RAV | 2834 × 2820 × 2828 | 1,016,813,441 | 2.42 |
| TOTAL FLOPS |  |  | 2.4 ZFLOPS |  |

<sup>a</sup>M, E, S, SH, and V models represent SARS-CoV-2 Delta strain. <sup>b</sup>Abbreviations used throughout document. <sup>c</sup>Periodic boundary dimensions. <sup>d</sup>Total number of atoms.
<sup>e</sup>Total aggregate simulation time, including heating and equilibration runs.
<sup>f</sup>Simulated with NAMD. <sup>g</sup>Simulated with NAMD, AMBER, and GROMACS.

#### 5.1.1 Simulating the SARS-CoV-2 structural proteins

Fully glycosylated Delta spike (S) structures in open and closed conformations were built based on WT constructs from Casalino et al. (Casalino et al., 2020) with the following mutations: T19R, T95I, G142D, E156G, Δ157-158, L452R, T478K, D614G, P681R, and D950N (Kannan et al.,2021, McCallum et al., 2021). Higher resolved regions were grafted from PDB 7JJI (Bangaru et al., 2020). Additionally, coordinates of residues 128-167 – accounting for a drastic conformational change seen in the Delta variant S – graciously made available to us by the Veesler Lab, were similarly grafted onto our constructs (McCallum et al., 2021). Finally, the S proteins were glycosylated following work by Casalino et al. (Casalino et al., 2020). By incorporating the Veesler Lab’s bleeding-edge structure (McCallum et al., 2021) and highly resolved regions from 7JJI (Bangaru et al., 2020), our models represent the most complete and accurate structures of the Delta S to date. The S proteins were inserted into membrane patches and equilibrated for 3 × 110 ns. For nonequilibrium and weighted ensemble simulations, a closed S head (SH, residues 13-1140) was constructed by removing the stalk from the full-length closed S structure, then resolvated, neutralized, minimized, and subsequently passed to WE and D-NEMD teams. The M protein was built from a structure graciously provided by the Feig Lab (paper in prep). The model was inserted into a membrane patch and equilibrated for 700 ns. RMSD-based clustering was used to select a stable starting M protein conformation. From the equilibrated and clustered M structure, VMD’s Mutator plugin (Humphrey et al., 1996) was used to incorporate the I82T mutation onto each M monomer to arrive at the Delta variant M. To construct the most complete E protein model to-date, the structure was patched together by resolving incomplete PDBs 5X29 (Surya et al., 2018), 7K3G (Mandala et al., 2020) and 7M4R (Chai et al., 2021). To do so, the transmembrane domain (residues 8-38) from 7K3G were aligned to the the N-terminal domain (residues 1-7) and residues 39 to 68 of 5X29 and residues 69 to 75 of 7M4R by their *C_α_* atoms. E was then inserted into a membrane patch and equilibrated for 40 ns.

#### 5.1.2 Constructing the SARS-CoV-2 Delta virion

The SARS-CoV-2 Delta virion (V) model was constructed following Casalino et al. (Casalino et al., 2021) using CHARMM-GUI (Lee et al., 2016), LipidWrapper (Durrant and Amaro, 2014), and Blender (Blender Online Community, 2020), using a 350 Å lipid bilayer with an equilibrium area per lipid of 63 Å^2^ and a 100 nm diameter Blender icospherical surface mesh (Turoňová et al., 2020). The resulting lipid membrane was solvated in a 1100 Å^3^ waterbox and subjected to 4 rounds of equilibration and patching (Casalino et al., 2021). 360 M dimers and 4 E pentamers were then tiled onto the surface, followed by random placement of 29 full-length S proteins (9 open, 20 closed) according to experimentally observed S protein density (Ke et al., 2020). M and E proteins were oriented with intravirion C-termini. After solvation in a 1460 Å waterbox, the complete V model tallied >305 million atoms (Table 1). V was equilibrated for 41 ns prior to placement in the respiratory aerosol (RA) model. The equilibrated membrane was 90 nm in diameter and remains in close structural agreement with the experimental studies (Ke et al., 2020).

#### 5.1.3 Building and simulating the respiratory aerosol

Respiratory aerosols contain a complex mixture of chemical and biological species. We constructed a respiratory aerosol (RA) fluid based on a composition from artificial saliva and surrogate deep lung fluid recipes (Walker et al., 2021). This recipe includes 0.7 mM DPPG, 6.5 mM DPPC, 0.3 mM cholesterol, 1.4 mM Ca^2+^, 0.8 mM Mg^2+^, and 142 mM Na^+^ (Vejerano and Marr, 2018, Walker et al., 2021), human serum albumin (ALB) protein, and a composition of mucins (Fig. 3). Mucins are long polymer-like structures that are decorated by dense, heterogeneous, and complex regions of O-glycans. This work represents the first of its kind as, due to their complexity, the O-glycosylated regions of mucins have never before been constructed for molecular simulations. Two short (m_1_, m_2_, ~5 nm) and three long (m_3_, m_4_, m_5_ ~55 nm) mucin models were constructed following known experimental compositions of protein and glycosylation sequences (Hughes et al., 2019, Mariethoz et al., 2018, Markovetz et al., 2019, Symmes et al., 2018, Thomsson et al., 2005) with ROSETTA (Raveh et al., 2010) and CHARMM GUI Glycan Modeller (Jo et al., 2011). Mucin models (short and long) were solvated, neutralized by charge matching with Ca^2+^ ions, minimized, and equilibrated for 15-25 ns each (Table 1). Human serum albumin (ALB), which is also found in respiratory aerosols, was constructed from PDB 1AO6 (Sugio et al., 1999). ALB was solvated, neutralized, minimized, and equilibrated for 7ns. Equilibrated structures of ALB and the three long mucins were used in construction of the RAV with m3+m4+m5 added at 6 g/mol and ALB at 4.4 g/mol.

**Figure 3:**
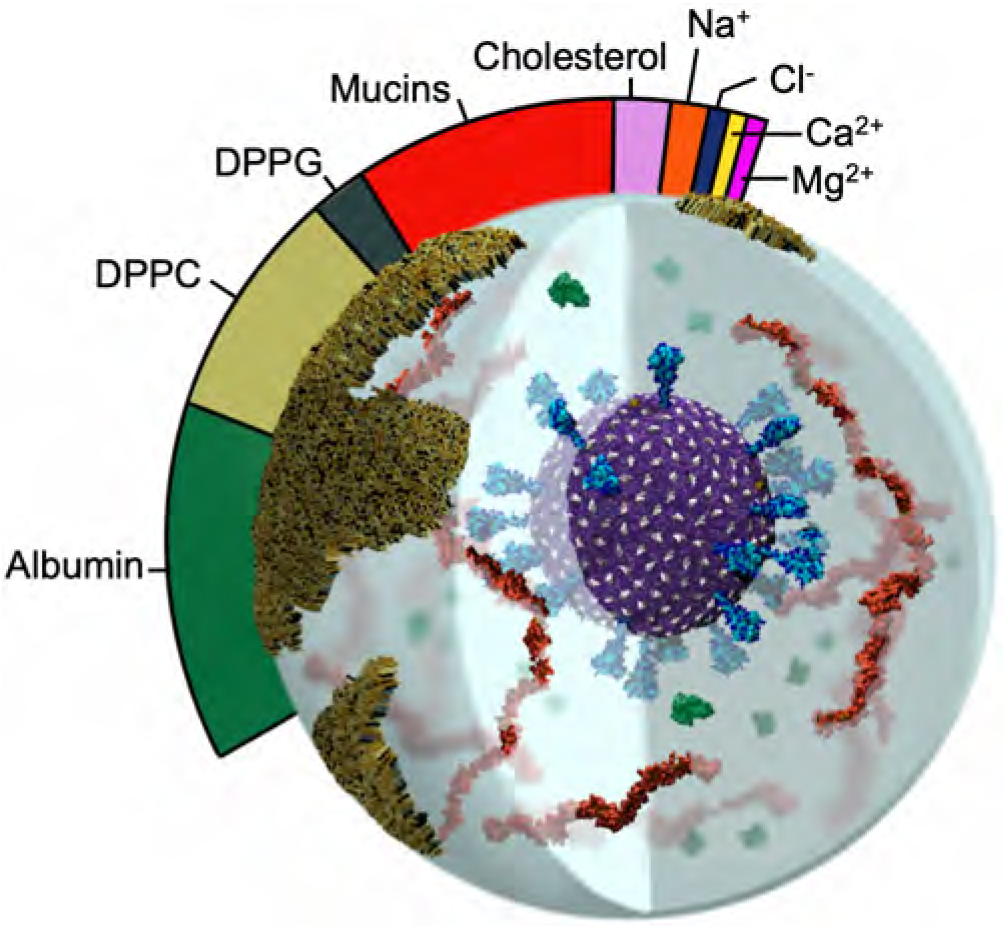
Image of RAV with relative mass ratios of RA molecular components represented in the colorbar. Water content is dependent on the relative humidity of the environment and is thus omitted from the molecular ratios.

#### 5.1.4 Constructing the respiratory aerosolized virion model

A 100 nm cubic box with the RA fluid recipe specified above was built with PACKMOL (Martínez et al., 2009), minimized, equilibrated briefly on TACC Frontera, then replicated to form a 300 nm cube. The RA box was then carved into a 270 nm diameter sphere. To make space for the placement of V within the RA, a spherical selection with volume corresponding to that of the V membrane + S crown (radius 734 Å) was deleted from the center of the RA. The final equilibrated V model, including surrounding equilibrated waters and ions (733 Å radius), was translated into the RA. Atom clashes were resolved using a 1.2 Å cutoff. Hydrogen mass repartitioning (Hopkins et al.,2015) was applied to the structure to improve performance. The simulation box was increased to 2800 Å per side to provide a 100 Å vacuum atmospheric buffer. The RAV simulation was conducted in an NVT ensemble with a 4 fs timestep. After minimizing, the RAV was heated to 298 K with 0.1 kcal/mol Å^2^ restraints on the viral lipid headgroups, then equilibrated for 1.5 ns. Finally, a cross-section of the RAV model – including and open S, m1/m2, and ALB (called the SMA system) – was constructed with PACKMOL to closely observe atomic scale interactions within the RAV model (Fig. 4).

**Figure 4:**
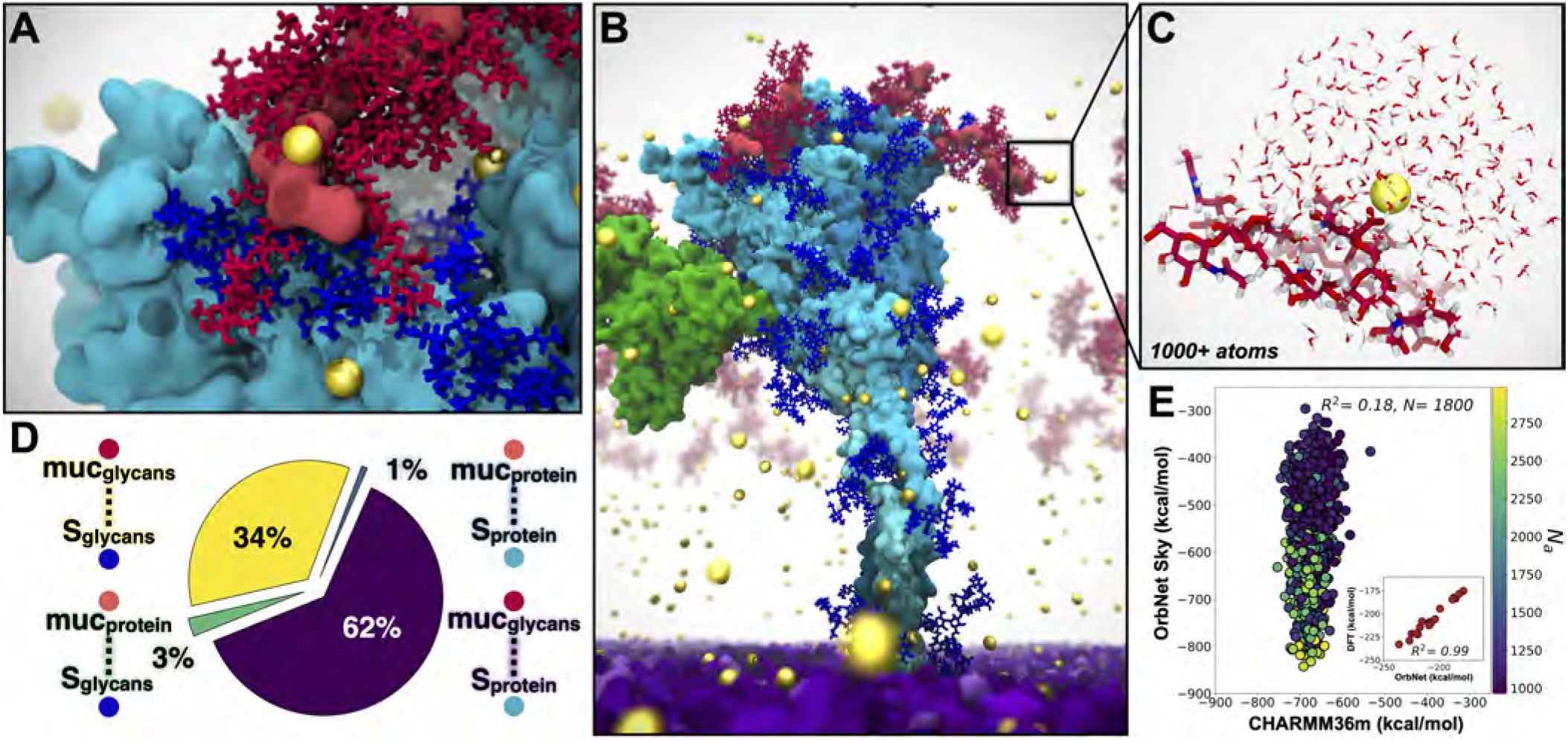
SMA system captured with multiscale modeling from classical MD to AI-enabled quantum mechanics. For all panels: S protein shown in cyan, S glycans in blue, m_1_/m_2_ shown in red, ALB in orange, Ca^2+^ in yellow spheres, viral membrane in purple. A) Interactions between mucins and S facilitated by glycans and Ca^2+^. B) Snapshot from SMA simulations. C) Example Ca^2+^ binding site from SMA simulations (1800 sites, each 1000+ atoms) used for AI-enabled quantum mechanical estimates from OrbNet Sky. D) Quantification of contacts between S and mucin from SMA simulations. E) OrbNet Sky energies vs CHARMM36m energies for each sub-selected system, colored by total number of atoms. Performance of OrbNet Sky vs. DFT in subplot (*ω*B97x-D3/def-TZVP, R^2^=0.99, for 17 systems of peptides chelating Ca^2+^ (Hu et al., 2021)). Visualized with VMD.

**Figure 5:**
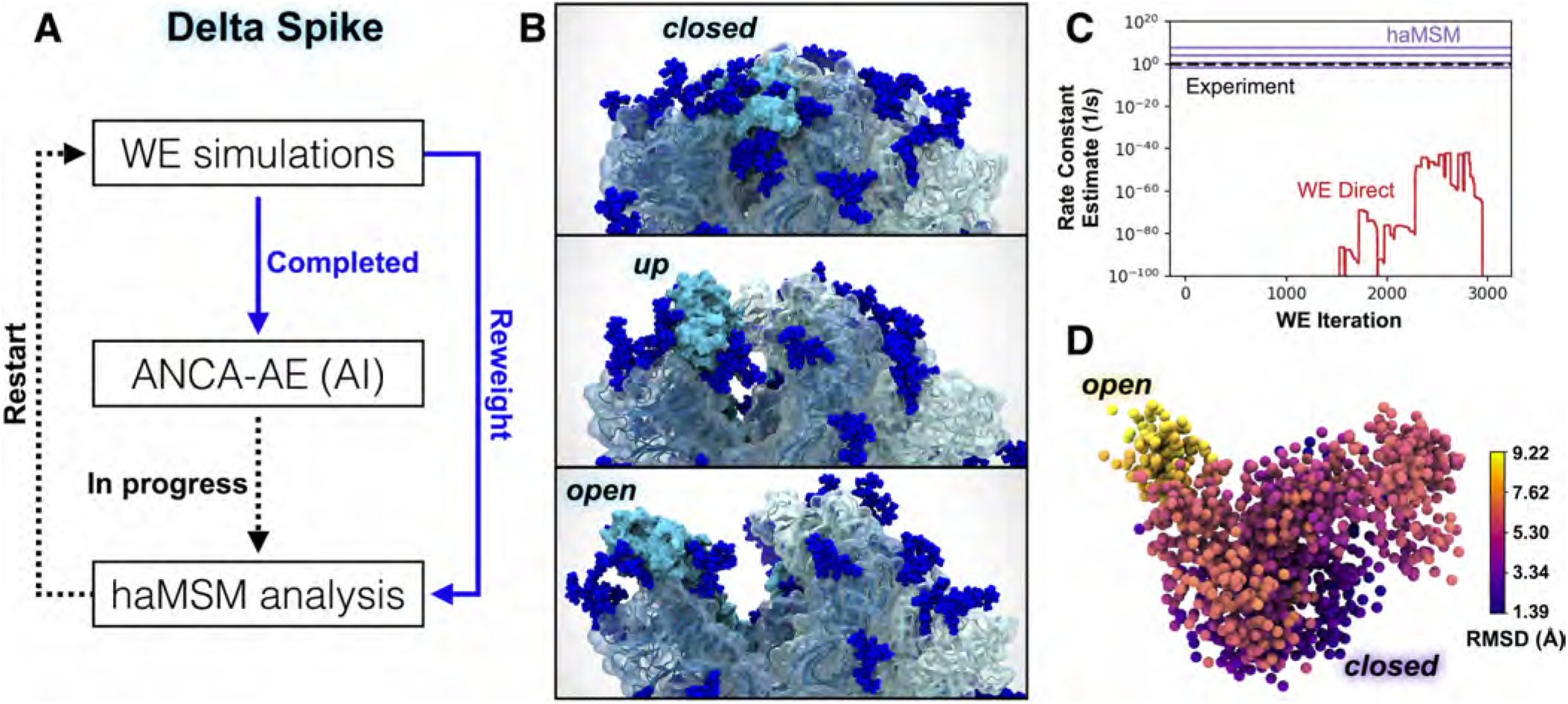
Delta-variant spike opening from WE simulations, and AI/haMSM analysis. A) The integrated workflow. B) Snapshots of the ‘down’, ‘up’, and ‘open’ states for Delta S-opening from a representative pathway generated by WE simulation, which represents ~10^5^speedup compared to conventional MD. C) Rate-constant estimation with haMSM analysis of WE data (purple lines) significantly improves direct WE computation (red), by comparison to experimental measurement (black dashed). Varying haMSM estimates result from different featurizations which will be individually cross-validated. D) The first three dimensions of the ANCA-AE embeddings depict a clear separation between the closed (darker purple) and open (yellow) conformations of the Delta spike. A sub-sampled landscape is shown here where each sphere represents a conformation from the WE simulations and colored with the root-mean squared deviations (Å) with respect to the closed state. Visualized with VMD.

**Figure 6:**
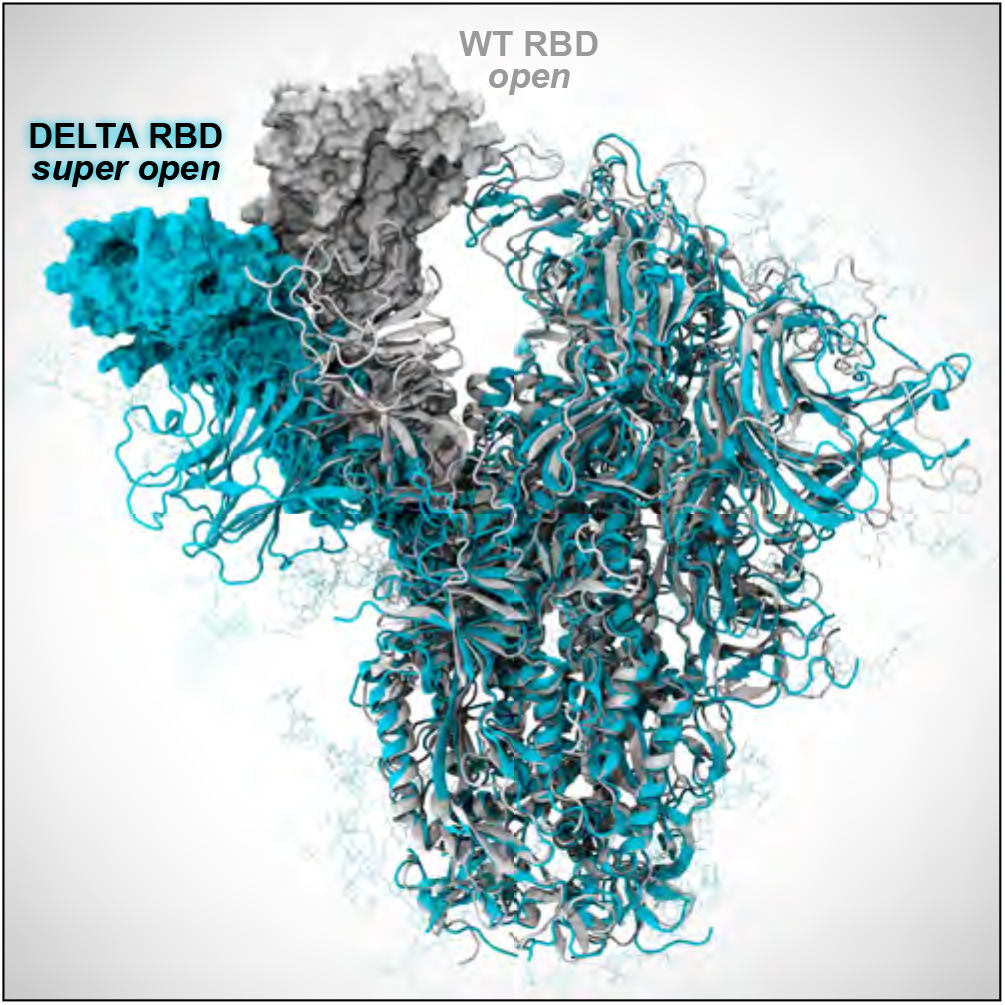
WE simulations reveal a dramatic opening of the Delta S (cyan), compared to WT S (white). While further investigation is needed, this super open state seen in the Delta S may indicate increased capacity for binding to human host-cell receptors.

### 5.2 Parameter evaluation with OrbNet

Comparison to quantum methods reveals significant polarization effects, and shows that there is opportunity to improve the accuracy of fixed charge force fields. For the large system sizes associated with solvated Ca^2+^-protein interaction motifs (over 1000 atoms, even in aggressively truncated systems) conventional quantum mechanics methods like density functional theory (DFT) are impractical for analyzing a statistically significant ensemble of distinct configurations (see discussion in Performance Results). In contrast, OrbNet allows for DFT accuracy with over 1000-fold speed-up, providing a useful method for benchmarking and refining the forcefield simulation parameters with quantum accuracy (Christensen et al., 2021). To confirm the accuracy of OrbNet versus DFT (*ω*B97X-D/def2-TZVP), the inset of Fig. 4E correlates the two methods for the Ca^2+^-binding energy in a benchmark dataset of small Ca^2+^-peptide complexes (Hu et al., 2021). The excellent correlation of OrbNet and DFT for the present use case is clear from the inset figure; six datapoints were removed from this plot on the basis of a diagnostic applied to the semi-empirical GFN-xTB solution used for feature generation of OrbNet (Christensen et al., 2021).

Fig. 4E presents a comparison of the validated OrbNet method with the CHARMM36m force field for 1800 snapshots taken from the SMA MD simulations. At each snapshot, a subsystem containing a solvated Ca^2+^-protein complex was extracted (Fig. 4E), with protein bonds capped by hydrogens. For both OrbNet and the force field, the Ca^2+^-binding energy was computed and shown in the correlation plot. Lack of correlation between OrbNet and the force field identifies important polarization effects, absent in a fixed charge description. Similarly, the steep slope of the best-fit line in Fig. 4E reflects the fact that some of the configurations sampled using MD with the CHARMM36m force field are relatively high in energy according to the more accurate OrbNet potential. This approach allows us to test and quantify limitations of empirical force fields, such as lack of electronic polarization.

The practicality of OrbNet for these simulation snapshots with 1000+ atoms offers a straightforward multiscale strategy for refining the accuracy of the CHARMM36m force field. By optimizing the partial charges and other force field parameters, improved correlation with OrbNet for the subtle Ca^2+^-protein interactions could be achieved, leading to near-quantum accuracy simulations with improved configurational sampling. The calculations presented here present a proof-of-concept of this iterative strategy.

### 5.3 AI-WE simulations of Delta spike opening

While our previous WE simulations of the WT SARS-CoV-2 S-opening (Sztain et al., 2021) were notable in generating pathways for a seconds-timescale process of a massive system, we have made two critical technological advancements in the WESTPA software that greatly enhance the efficiency and analysis of WE simulations. These advances enabled striking observations of Delta-variant S opening (Figs. 5 and 6). First, in contrast to prior manual bins for controlling trajectory replication, we have developed automated and adaptive binning that enables more efficient surmounting of large barriers via early identification of “bottleneck” regions (Torrillo et al., 2021). Second, we have parallelized, memory-optimized, and implemented data streaming for the history-augmented Markov state model (haMSM) analysis scheme (Copperman and Zuckerman,2020) to enable application to the TB-scale S-opening datasets. The haMSM approach estimates rate constants from simulations that have not yet reached a steady state (Suarez et al., 2014).

Our WE simulations generated >800 atomically detailed, Delta-variant S-opening pathways (Figs. 5B and 6) of the receptor binding domain (RBD) switching from a glycan-shielded ‘down’ to an exposed ‘up’ state using 72 *μ*s of total simulation time within 14 days using 192 NVIDIA V100 GPUs at a time on TACC’s Longhorn supercomputer. Among these pathways, 83 reach an ‘open’ state that aligns with the structure of the human ACE2-bound WT S protein (Benton et al., 2020) and 18 reach a dramatically open state (Fig. 6). Our haMSM analysis of WT WE simulations successfully provided long-timescale (steady-state) rate constants for S-opening based on highly transient information (Fig. 5C).

We also leveraged a simple, yet powerful unsupervised deep learning method called Anharmonic Conformational Analysis enabled Autoencoders (ANCA-AE) (Clyde et al., 2021) to extract conformational states from our long-timescale WE simulations of Delta spike opening (Fig. 5A,D). ANCA-AE first minimizes the fourth order correlations in atomistic fluctuations from MD simulation datasets and projects the data onto a low dimensional space where one can visualize the anharmonic conformational fluctuations. These projections are then input to an autoencoder that further minimizes non-linear correlations in the atomistic fluctuations to learn an embedding where conformations are automatically clustered based on their structural and energetic similarity. A visualization of the first three dimensions from the latent space articulates the RBD opening motion from its closed state (Fig. 5D). It is notable that while other deep learning techniques need special purpose hardware (such as GPUs), the ANCA-AE approach can be run with relatively modest CPU resources and can therefore scale to much larger systems (e.g., the virion within aerosol) when optimized.

### 5.4 D-NEMD explores pH effects on Delta spike

We performed D-NEMD simulations of the SH system with GRO-MACS (Abraham et al., 2015) using a ΔpH=2.0 (from 7.0 to 5.0) as the external perturbation. We ran 3 200-ns equilibrium MD simulations of SH to generate 87 configurations (29 configurations per replicate) that were used as the starting points for multiple short (10 ns) D-NEMD trajectories under the effect of the external perturbation (ΔpH=2.0). The effect of a ΔpH was modelled by changing the protonation state of histidines 66, 69, 146, 245, 625, 655, 1064, 1083, 1088, and 1101 (we note that other residues may also become protonated (Lobo and Warwicker, 2021); the D-NEMD approach can also be applied to examine those). The structural response of the S to the pH decrease was investigated by measuring the difference in the position for each *Cα* atom between the equilibrium and corresponding D-NEMD simulation at equivalent points in time (Oliveira et al., 2021a), namely after 0, 0.1, 1, 5 and 10 ns of simulation. The D-NEMD simulations reveal that pH changes, of the type expected in aerosols, affect the dynamics of functionally important regions of the spike, with potential implications for viral behavior (Fig. 7). As this approach involves multiple short independent non-equilibrium trajectories, it is well suited for cloud computing. All D-NEMD simulations were performed using Oracle Cloud.

**Figure 7:**
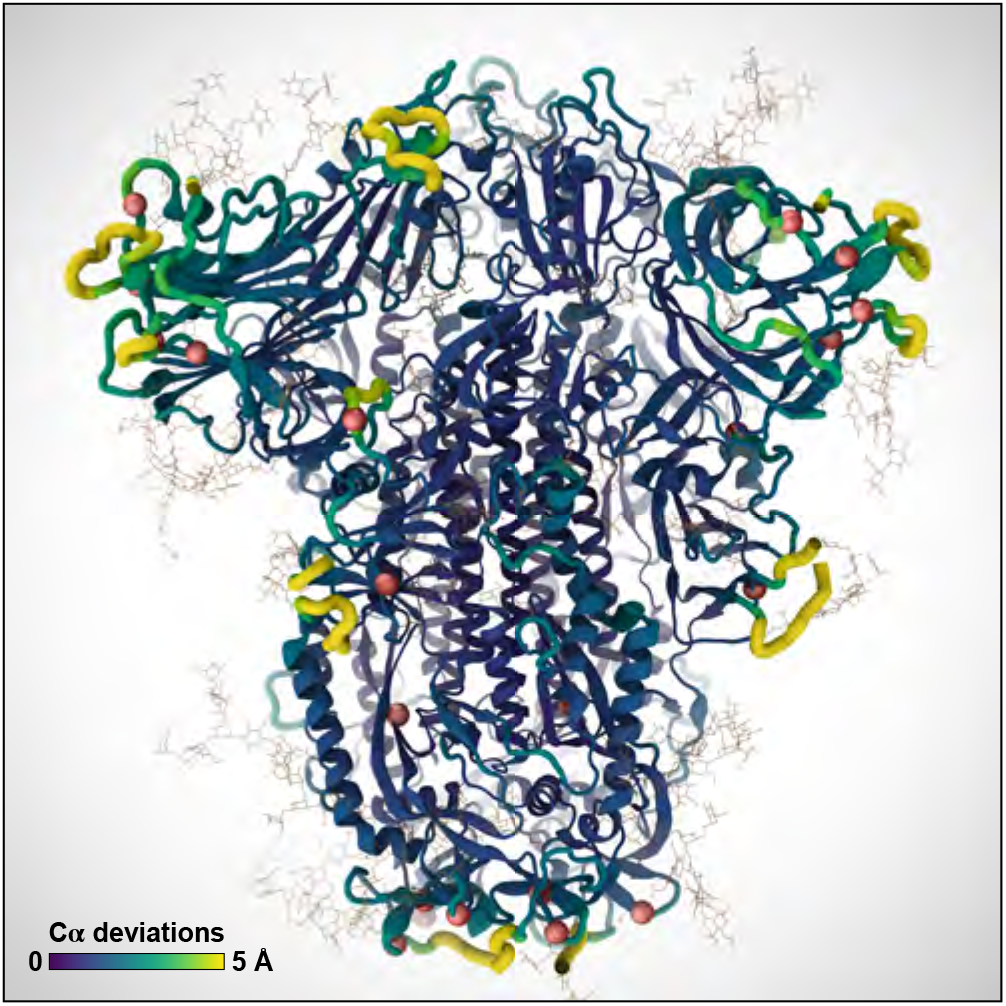
D-NEMD simulations reveal changes in key functional regions of the S protein, including the receptor binding domain, as the result of a pH decrease. Color scale and ribbon thickness indicate the degree of deviation of C*α* atoms from their equilibrium position. Red spheres indicate the location of positively charged histidines.

## 6 HOW PERFORMANCE WAS MEASURED

### 6.1 WESTPA

For the WE simulations of spike opening using WESTPA, we defined the time to solution as the total simulation time required to generate the first spike opening event. Spike opening is essentially impossible to observe via conventional MD. WESTPA simulations were run using the AMBER20 dynamics engine and 192 NVIDIA V100 GPUs at a time on TACC’s Longhorn supercomputer.

### 6.2 NAMD

NAMD performance metrics were collected using hardware performance counters for FLOPs/step measurements, and application-internal timers for overall simulation rates achieved by production runs including all I/O for simulation trajectory and checkpoint output. NAMD FLOPs/step measurements were conducted on TACC Frontera, by querying hardware performance counters with the `rdmsr` utility from Intel msr-tools^1^ and the “TACC stats” system programs.^2^ For each simulation, FLOP counts were measured for NAMD simulation runs of two different step counts. The results of the two simulation lengths were subtracted to eliminate NAMD startup operations, yielding an accurate estimate of the marginal FLOPs per step for a continuing simulation (Phillips et al., 2002). Using the FLOPs/step values computed for each simulation, overall FLOP rates were computed by dividing the FLOPs/step value by seconds/step performance data reported by NAMD internal application timers during production runs.

**Table 2:** MD simulation floating point ops per timestep.

| MD Simulation | Code | Atoms | <sup>a</sup>FLOPs/step |
| --- | --- | --- | --- |
| Spike, head | AMBER, GROMACS | 0.6M | 62.14 GFLOPs/step |
| Spike | NAMD | 1.7M | 43.05 GFLOPs/step |
| S+m<sub>1</sub>/m<sub>2</sub>+ALB | NAMD | 2.1M | 54.86 GFLOPs/step |
| Resp. Aero.+Vir. | NAMD | 1B | 25.81 TFLOPs/step |

<sup>a</sup> FLOPs/step data were computed by direct FLOP measurements from hardware performance counters for NAMD simulations, or by using the application-reported FLOP rates and ns/day simulation performance in the case of GROMACS.

### 6.3 GROMACS

GROMACS 2020.4 benchmarking was performed on Oracle Cloud Infrastructure (OCI)^3^ compute shape BM.GPU4.8 consisting of 8×NVIDIA A100 tensor core GPUs, and 64 AMD Rome CPU cores. The simulation used for benchmarking contained 615,563 atoms and was run for 500,000 steps with 2 fs time steps. The simulations were run on increasing numbers of GPUs, from 1 to 8, using 8 CPU cores per GPU, running for both the production (Nose-Hoover) and GPU-accelerated (velocity rescaling) thermostats. Particle–mesh Ewald (PME) calculations were pinned to a single GPU, with additional GPUs for multi-GPU jobs used for particle–particle calculations. Performance data (ns/day and average single-precision TFLOPS, calculated as total number of TFLOPs divided by total job walltime) were reported by GROMACS itself. Each simulation was repeated four times and average performance figures reported.

## 7 PERFORMANCE RESULTS

### 7.1 NAMD performance

NAMD was used to perform all of the simulations listed in Table 1, except for the closed spike “SH” simulations described further below. With the exception of the aerosol and virion simulation, the other NAMD simulations used conventional protocols and have performance and parallel scaling characteristics that closely match the results reported in our previous SARS-CoV-2 research (Casalino et al., 2021). NAMD 2.14 scaling performance for the one billion-atom respiratory aerosol and virion simulation run on ORNL Summit is summarized in Tables 3 and 4. A significant performance challenge associated with the aerosol virion simulation relates to the roughly 50% reduction in particle density as compared with a more conventional simulation with a fully populated periodic cell. The reduced particle density results in large regions of empty space that nevertheless incur additional overheads associated with both force calculations and integration, and creates problems for the standard NAMD load balancing scheme that estimates the work associated with the cubic “patches” used for parallel domain decomposition. The PME electrostatics algorithm and associated 3-D FFT and transpose operations encompass the entire simulation unit cell and associated patches, requiring involvement in communication and reduction operations despite the inclusion of empty space. Enabling NAMD diagnostic output on a 512-node 1B-atom aerosol and virion simulation revealed that ranks assigned empty regions of the periodic cell had 66 times the number of fixed-size patches as ranks assigned dense regions. The initial load estimate for an empty patch was changed from a fixed 10 atoms to a runtime parameter with a default of 40 atoms, which reduced the patch ratio from 66 to 19 and doubled performance on 512 nodes.

**Table 3:** NAMD performance: Respiratory Aerosol + Virion, 1B atoms, 4 fs timestep w/ HMR, and PME every 3 steps.

| Nodes | Summit<br>CPU + GPU | Speedup | Efficiency |
| --- | --- | --- | --- |
| 256 | 4.18 ns/day | ~1.0× | ~100% |
| 512 | 7.68 ns/day | 1.84× | 92% |
| 1024 | 13.64 ns/day | 3.27× | 81% |
| 2048 | 23.10 ns/day | 5.53× | 69% |
| 4096 | 34.21 ns/day | 8.19× | 51% |

**Table 4:** Peak NAMD FLOP rates, ORNL Summit.

| NAMD Simulation | Atoms | Nodes | Sim rate | Performance |
| --- | --- | --- | --- | --- |
| Resp. Aero.+Vir. | 1B | 4096 | 34.21 ns/day | 2.55 PFLOPS |

### 7.2 WESTPA performance

Our time to solution for WE simulations of spike opening (to the “up” state) (Fig. 5) using the WESTPA software and AMBER20 was 14 *μs* of total simulation time, which was completed in 4 days using 192 NVIDIA V100 GPUs at a time on TACC’s Longhorn supercomputer. For reference, conventional MD would require an expected ~5 orders of magnitude more computing. The WESTPA software is highly scalable, with nearly perfect scaling out to >1000 NVIDIA V100 GPUs and this scaling is expected to continue until the filesystem is saturated. Thus, WESTPA makes optimal use of large supercomputers and is limited by filesystem I/O due to the periodic restarting of trajectories after short time intervals.

### 7.3 AI-enhanced WE simulations

DeepDriveMD is a framework to coordinate the concurrent execution of ensemble simulations and drive them using AI models (Brace et al., 2021a, Lee et al., 2019). DeepDriveMD has been shown to improve the scientific performance of diverse problems: from protein folding to conformation of protein-ligand complexes. We coupled WESTPA to DeepDriveMD, which is responsible for resource dynamism and concurrent heterogeneous task execution (ML and AMBER). The coupled workflow was executed on 1024 nodes on Summit (OLCF), and, in spite of the spatio-temporal heterogeneity of tasks involved, the resource utilization was in the high 90%. Consistent with earlier studies, the coupling of WESTPA to DeepDriveMD results in a 100x improvement in the exploration of phase space.

### 7.4 GROMACS performance

Figure 8 shows GROMACS parallelizes well across the 8 NVIDIA A100 GPUs available on each BM.GPU4.8 instance used in the *Cluster in the Cloud*^4^ running on OCI. There is a performance drop for two GPUs due to inefficient division of the PME and particle-particle tasks. Methods to address this exist for the two GPU case (Páll et al.,2020), but were not adopted as we were targeting maximum raw performance across all 8 GPUs. Production simulations achieved 27% of the peak TFLOPS available from the GPUs. Multiple simulations were run across 10 such compute nodes, enabling the ensemble to run at an average combined speed of 425 TFLOPS and sampling up to 1μs/day. We note that the calculations will be able to run 20%–40% faster once the Nose-Hoover thermostat that is required for the simulation is ported to run on the GPU. Benchmarking using a velocity rescaling thermostat that has been ported to GPU shows that this would enable the simulation to extract 34% of the peak TFLOPS from the cards, enabling each node to achieve an average speed of 53.4 TFLOPS, and 125 ns/day. A cluster of 10 nodes would enable GROMACS to run at an average combined speed of over 0.5 PFLOPs, simulating over 1.2 μs/day.

**Figure 8:**
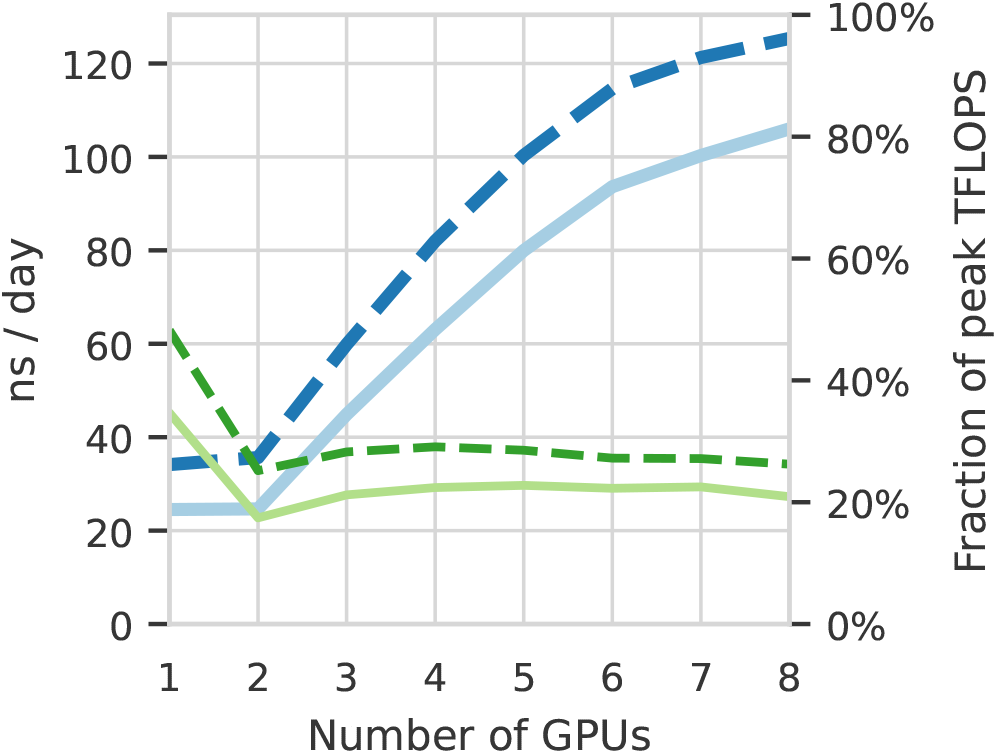
GROMACS performance across 1–8 A100 GPUs in ns/day (thicker, blue lines) and the fraction of maximum theoretical TFLOPS (thinner, green lines); production setup shown with solid line, and runs with the GPU-accelerated thermostat in dashed.

A significant innovation is that this power is available on demand: Cluster in the Cloud with GPU-optimized GROMACS was provisioned and benchmarked within one day of inception of the project. This was handed to the researcher, who submitted the simulations. Automatically, up to ten BM.GPU4.8 compute nodes were provisioned on-demand based on requests from the Slurm scheduler. These simulations were performed on OCI, using *Cluster in the Cloud* (Williams, 2021) to manage automatic scaling.

Cluster in the Cloud was configured to dynamically provision and terminate computing nodes based on the workload. Simulations were conducted using GROMACS 2020.4 compiled with CUDA support. Multiple simultaneous simulations were conducted, with each simulation utilizing a single BM.GPU4.8 node without multinode parallelism.

This allowed all production simulations to be completed within 2 days. The actual compute cost of the project was less than $6125 USD (on-demand OCI list price). The huge reduction in “time to science” that low-cost cloud enables changes the way that researchers can access and use HPC facilities. In our opinion, such a setup enables “exclusive on-demand” HPC capabilities for the scientific community for rapid advancement in science.

### 7.5 OrbNet performance

Prior benchmarking reveals that OrbNet provides over 1000-fold speedup compared to DFT (Christensen et al., 2021). For the calculations presented here, the cost of corresponding high quality range-separated DFT calculations (wB97X-D/def2-TZVP) can be estimated. In Fig. 4E, we consider system sizes which would require 14,000–47,000 atomic orbitals for wB97X-D/def2-TZVP, exceeding the range of typical DFT evaluations. Estimation of the DFT computational cost of the 1811 configurations studied in Fig. 4E suggests a total of 115M core-hours on NERSC Cori Haswell nodes; in contrast, the OrbNet calculations for the current study require only 100k core-hours on the same nodes. DFT cost estimates were based on extrapolation from a dataset of over 1M ChEMBL molecules ranging in size from 40 to 107 atom systems considering only the cubic cost component of DFT (Christensen et al., 2021).

## 8 IMPLICATIONS

Our major scientific achievements are:

1. We showcase an extensible AI-enabled multiscale computational framework that bridges time and length scales from electronic structure through whole aerosol particle morphology and dynamics.
2. We develop all-atom simulations of respiratory mucins, and use these to understand the structural basis of interaction with the SARS-CoV-2 spike protein. This has implications for viral binding in the deep lung, which is coated with mucins. We expect the impact of our mucin simulations to be far reaching, as malfunctions in mucin secretion and folding have been implicated in progression of severe diseases such as cancer and cystic fibrosis.
3. We present a significantly enhanced all-atom model and simulation of the SARS-CoV-2 Delta virion, which includes the hundreds of tiled M-protein dimers and the E-protein ion channels. This model can be used as a basis to understand why the Delta virus is so much more infectious than the WT or alpha variants.
4. We develop an ultra-large (1 billion+) all-atom simulation capturing massive chemical and biological complexity within a respiratory aerosol. This simulation provides the first atomic level views of virus-laden aerosols and is already serving as a basis to develop an untold number of experimentally testable hypotheses. An immediate example suggests a mechanism through which mucins and other species, e.g., lipids, which are present in the aerosol, arrange to protect the molecular structure of the virus, which otherwise would be exposed to the air-water interface. This work also opens the door for developing simulations of other aerosols, e.g., sea spray aerosols, that are involved in regulating climate.
5. We evidence how changes in pH, which are expected in the aerosol environment, may alter dynamics and allosteric communication pathways in key functional regions of the Delta spike protein.
6. We characterize atomically detailed pathways for the spike-opening process of the Delta variant using WE simulations, revealing a dramatically open state that may facilitate binding to human host cells.
7. We demonstrate how parallelized haMSM analysis of WE data can provide physical rate estimates of spike opening, improving prior estimates by many orders of magnitude. The pipeline can readily be applied to the any variant spike protein or other complex systems of interest.
8. We show how HPC and cloud resources can be used to significantly drive down time-to-solution for major scientific efforts as well as connect researchers and greatly enable complex collaborative interactions.
9. We demonstrate how AI coupled to HPC at multiple levels can result in significantly improved effective performance, e.g., with AI-driven WESTPA, and extend the reach and domain of applicability of tools ordinarily restricted to smaller, less complex systems, e.g., with OrbNet.
10. While our work provides a successful use case, it also exposes weaknesses in the HPC ecosystem in terms of support for key steps in large/complex computational science campaigns. We find lack of widespread support for high performance remote visualization and interactive graphical sessions for system preparation, debugging, and analysis with diverse science tools to be a limiting factor in such efforts.

## ACKNOWLEDGMENTS

We thank Prof. Kim Prather for inspiring and informative discussions about aerosols and for her commitment to convey the airborne nature of SARS-CoV-2. We thank D. Veesler for sharing the Delta spike NTD coordinates in advance of publication. We thank B. Messer, D. Maxwell, and the Oak Ridge Leadership Computing Facility at Oak Ridge National Laboratory supported by the DOE under Contract DE-AC05-00OR22725. We thank the Texas Advanced Computing Center Frontera team, especially D. Stanzione and T. Cockerill, and for compute time made available through a Director’s Discretionary Allocation (NSF OAC-1818253). We thank the Argonne Leadership Computing Facility supported by the DOE under DE-AC02-06CH11357. We thank the Pittsburgh Supercomputer Center for providing priority queues on Bridges-2 through the XSEDE allocation NSF TG-CHE060063. We thank N. Kern and J. Lee of the CHARMM-GUI support team for help converting topologies between NAMD and GROMACS. We thank J. Copperman, G. Simpson, D. Aristoff, and J. Leung for valuable discussions and support from NIH grant GM115805. NAMD and VMD are funded by NIH P41-GM104601. This work was supported by the NSF Center for Aerosol Impacts on Chemistry of the Environment (CAICE), National Science Foundation Center for Chemical Innovation (NSF CHE-1801971), as well as NIH GM132826, NSF RAPID MCB-2032054, an award from the RCSA Research Corp., a UC San Diego Moore’s Cancer Center 2020 SARS-CoV-2 seed grant, to R.E.A. This work was also supported by Oracle Cloud credits and related resources provided by the Oracle for Research program.

## Footnotes

1 https://github.com/intel/msr-tools

2 https://github.com/TACC/tacc_stats

3 https://www.oracle.com/cloud/

4 https://cluster-in-the-cloud.readthedocs.io/

